# Genetic Analysis Suggests a Surface of PAT-4 (ILK) that Interacts with UNC-112 (kindlin)

**DOI:** 10.1101/2022.03.18.484907

**Authors:** Hiroshi Qadota, Annie McPherson, Rachel Corbitt, Evan Kelton Dackowski, Yohei Matsunaga, Andres Oberhauser, Guy M. Benian

**Author notes:** Corresponding author: Guy M. Benian.

## Abstract

The transmembrane protein integrin plays a crucial role in the attachment of cells to the extracellular matrix. Integrin recruits many proteins intracellularly, including a four protein complex (Kindlin, Integrin linked kinase (ILK), PINCH, and parvin). *C. elegans* muscle provides an excellent model to study integrin adhesion complexes. In *C. elegans*, UNC-112 (Kindlin) binds to the cytoplasmic tail of PAT-3 (β-integrin) and to PAT-4 (ILK). We previously reported that PAT-4 binding to UNC-112 is essential for the binding of UNC-112 to PAT-3. Although there are crystal structures for ILK and a kindlin, there is no co-crystal structure available. To understand the molecular interaction between PAT-4 (ILK) and UNC-112 (Kindlin), we took a genetic approach. First, we isolated mutant PAT-4 proteins that cannot bind to UNC-112. Then, we isolated suppressor mutant UNC-112 proteins that restore interaction with mutant PAT-4 proteins. Second, these mutant PAT-4 proteins cannot localize to attachment structures in nematode muscle, but upon co-expression of an UNC-112 suppressor mutant protein, the mutant PAT-4 protein could localize to attachment structures. Third, overexpression of a mutant PAT-4 protein results in disorganization of the adhesion plaques at muscle cell boundaries, and co-expression of the UNC-112 supressor mutant protein alleviates this defect. Thus, we demonstrate that UNC-112 binding to PAT-4 is required for the localization and function of PAT-4 in integrin adhesion complexes in vivo. The missense mutations were mapped onto homology models of PAT-4 and UNC-112, and taking into account previously isolated mutations, we suggest a surface of PAT-4 that binds to UNC-112.

## Introduction

*Caenorhabditis elegans* muscle used for locomotion is located in the body wall and consists of 95 spindle-shaped mononuclear cells arranged in interlocking pairs that run the length of the animal in four quadrants. The myofibrils are restricted to a narrow ∼1.5-μm zone adjacent to the cell membrane along the outer side of the muscle cell. The thin filaments are attached to dense bodies (Z-disk analogs), and the center of the bundle of thick filaments (A-bands) are attached to M-lines. Moreover, all of the dense bodies and M-lines are anchored to the muscle cell membrane and extracellular matrix, which is attached to the hypodermis and cuticle. This allows the force of muscle contraction to be transmitted directly to the cuticle and allows movement of the whole animal (Waterston 1988; Moerman and Fire 1997; Moerman and Williams 2006; Gieseler et al. 2017). Thus, the nematode muscle M-lines and dense bodies serve the function of analogous sarcomeric structures in vertebrate muscle, and, in addition, because these structures are attached to the cell membrane and consist of integrin and integrin-associated proteins (see below), they are also similar to costameres of vertebrate muscle and focal adhesions of non-muscle cells. The membrane-proximal regions of the dense bodies and M-lines consist of integrin and associated proteins (Moerman and Williams 2006; Gieseler et al. 2017). Integrin-associated proteins are also used at the adhesion plaques that lie between adjacent muscle cells, and are likely involved in transmitting forces laterally (Moody et al. 2020). The cytoplasmic tail of β-integrin (PAT-3) is associated with four conserved proteins: UNC-112 (kindlin), PAT-4 (integrin-linked kinase, ILK), PAT-6 (α-parvin), and UNC-97 (PINCH). The cytoplasmic tail of PAT-3 binds to UNC-112, and UNC-112 binds to PAT-4 (Qadota et al. 2012; Mackinnon et al. 2002). PAT-4 also has been shown to interact with PAT-6 and UNC-97 (Mackinnon et al. 2002; Lin et al. 2003; Norman et al. 2007). Each of these proteins is required for myofibril assembly in embryonic and adult muscle (Williams and Waterston 1994; Hobert et al. 1999). All five proteins are localized to integrin adhesion sites (M-lines, dense bodies, and muscle cell boundaries) in muscle cells (Rogalski et al. 2000; Mackinnon et al. 2002; Lin et al. 2003; Hobert et al. 1999; Norman et al. 2007; Qadota et al. 2017). When UNC-112 was immunoprecipitated, the other 3 proteins (PAT-4, PAT-6, and UNC-97) were co-precipited, consistent with a four-protein complex in vivo (Qadota et al. 2014).

We have suggested a model for the assembly of integrin-associated proteins previously (Qadota et al. 2012): 1) PAT-4 binding to UNC-112 changes the conformation of UNC-112 from closed (inactive) to open (active), 2) the UNC-112-PAT-4 complex binds to the cytoplasmic tail of β-integrin, 3) PAT-4 recruits PAT-6 and UNC-97. Within this integrin-integrin associated protein complex, the UNC-112 / PAT-4 complex functions as a core. Previously, we reported mutations in UNC-112 that cannot bind to PAT-4 (Qadota et al. 2012), and its extragenic suppressor mutations in PAT-4 (Qadota et al. 2014). These mutations are useful for understanding the interaction surface between UNC-112 and PAT-4 (Qadota and Benian 2014), especially because, although crystal structures for an ILK and a kindlin have been reported, to date, there is no kindlin/ILK (or UNC-112/PAT-4) co-crystal structure. To obtain more molecular information about the UNC-112/PAT-4 interaction, in this study, we screened for and obtained mutant PAT-4 proteins that cannot bind to UNC-112, and then isolated mutant UNC-112 proteins that can restore the interaction. We expressed mutant PAT-4 and suppressor mutant UNC-112 proteins using a heat shock promoter. After transient expression (2 hours of heat shock), we examined muscle localization of mutant PAT-4 with and without suppressor UNC-112. After sustained expression (24 hours of heat shock) of these mutant proteins, we examined the organization of muscle adhesion complexes.

## Methods & Materials

### Screening for PAT-4 mutants that cannot bind to UNC-112

Mutations in PAT-4 were introduced randomly by error-prone PCR. We PCR-amplified full-length PAT-4 cDNA by an error-prone method (Qadota et al. 2012) using the following primers: 5′ (GAA GAT ACC CCA CCA AAC) and 3′ (AAA GAA GGC AAA ACG ATG), and a PAT-4 cDNA plasmid (pDM#280) (Qadota et al. 2012) as a template. These primers contain 30 base pairs of sequence that overlap regions in the acceptor plasmid. Cloning of error-prone PCR-amplified fragments into the acceptor plasmid was performed by exploiting yeast recombination *in vivo* (Takita et al. 1997). The mixture of the amplified PCR fragments (1 μg) and the acceptor plasmid (pACT; 1 μg) digested with XhoI was transformed into PJ69-4A harboring pGBDU-PAT-6. Transformed yeast cells were spread onto -Leu-Ura-His plates containing 2 mM 3-amino-1,2,4-triazole for identifying His^+^ colonies. His^+^ selection ensured that the mutagenized PAT-4 could still interact with PAT-6. This step was essential for eliminating clones with premature stop mutations or with many other mutations. His^+^ colonies were streaked onto an -Ade plate and screened for His^+^Ade^+^ colonies. After streaking on a 5-fluoroorotic acid plate to eliminate the URA3 marker bait plasmid (pGBDU-PAT-6), prey clones were isolated from yeast and amplified in *E. coli*. From a total of 48 His^+^Ade^+^ yeast colonies, 47 mutagenized clones were isolated. These prey clones were transformed separately into PJ69–4A carrying either pGBDU-PAT-6 or pGBDU-UNC-112 (full-length) to check for interaction with PAT-6 and interaction with UNC-112. Among 47 mutagenized PAT-4 clones, all 47 showed binding to PAT-6, but 16 of these clones could not bind to full-length UNC-112. DNA sequencing of these clones revealed the mutations (Table 1).

**Table 1.**
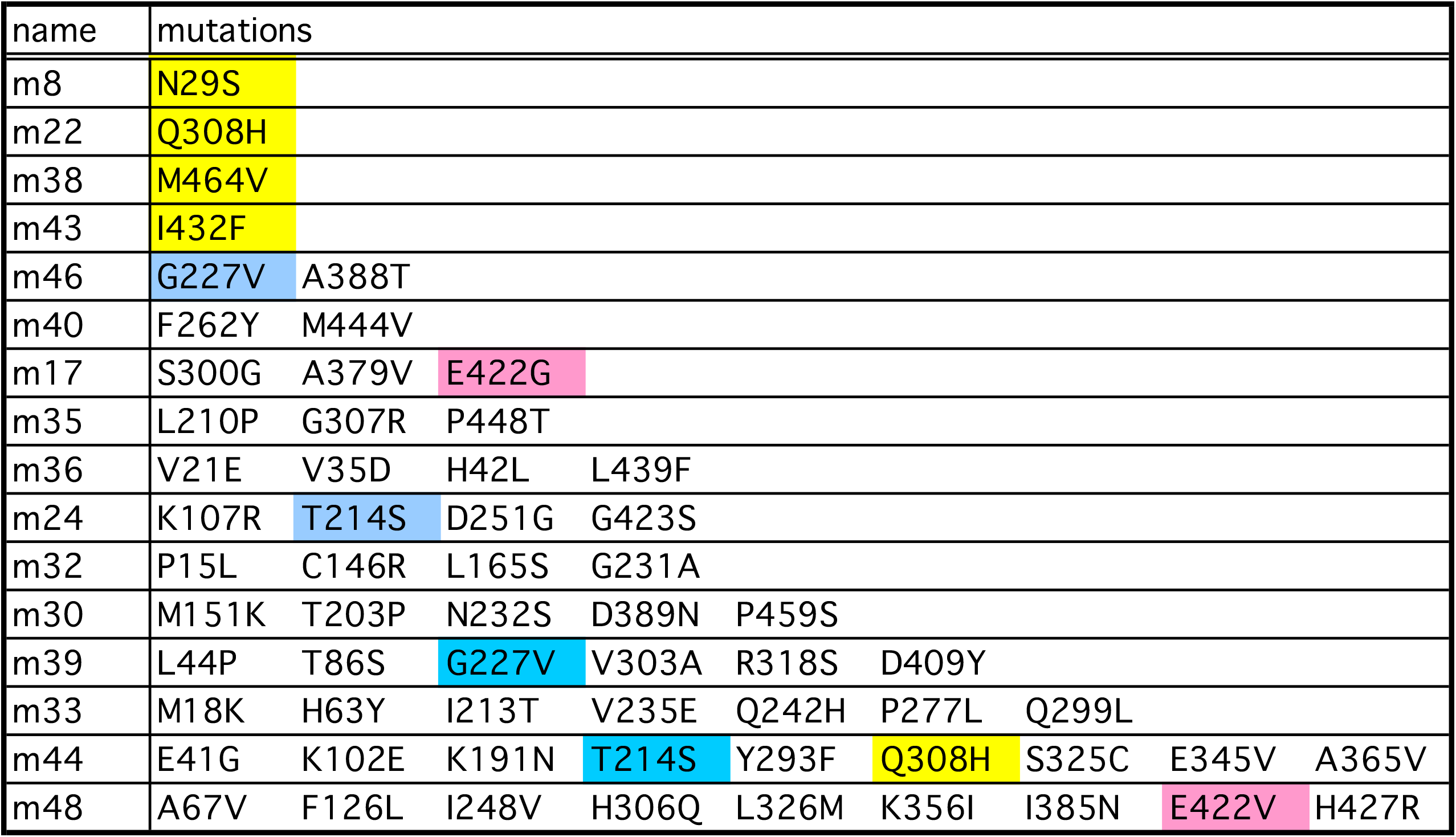
PAT-4 mutants that cannot bind to UNC-112. Yellow represents single mutations. Blue represents amino acid changes found in two independent clones. Pink represents residues mutated in two independent clones, but those changes result in different amino acid substitutions.

### Expression of HA-tagged PAT-4 wild type and mutants in *C. elegans* using heat shock promoters

Full-length PAT-4 cDNA without mutations, or with mutations Q308H, and I432F, were amplified by PCR using primers: EcoRV-PAT-4 (GCG GAT ATC ATG TCT TTG TCG ACT CAT TAC) and PAT-4-XhoI (CGC CTC GAG TCA TAA TAT CAT TCT CTC TAA) and cloned into the HA-tag vector, pKS-HA(Nhex2). From these pKS-HA (Nhex2) clones, NheI fragments (containing HA-tagged PAT-4 cDNAs) were cloned into the NheI site of *C. elegans* expression vectors, pPD49.78 and pPD49.83 (gifts from Dr. Andrew Fire, Stanford University). These vectors contain two different heat shock promoters. pPD49.78/83-HA-PAT-4 (WT, Q308H, and I432F) were mixed with pTG96 (SUR-5::NLS::GFP) as a transformation marker (Yochem et al. 1998) and injected into wild type N2 worms. Transgenic lines with extrachromosomal arrays containing pPD49.78/83-HA-PAT-4 (WT (called *sfEx56*), Q308H (called *sfEx54*), and I432F (called *sfEx55*), and pTG96 were established by picking GFP-positive worms using a GFP dissection microscope. Expression of the HA-tagged PAT-4 proteins (WT, Q308H, and I432F) was induced by incubation of the transgenic worms at 30 °C for 2 h (heat shock). We prepared worm lysates (Hannak et al. 2002) from transgenic worms with or without heat shock and examined the expression of HA-tagged PAT-4 proteins by Western blot, reacting with anti-HA (Sigma-Aldrich H3663; 1:200 dilution).

### Screening for UNC-112 mutants that restore binding to PAT-4 Q308H or PAT-4 I432F

We utilized random mutagenesis by PCR, as described previously (Qadota et al. 2014). UNC-112 cDNAs with mutations were amplified with two primers (U112-P1 (GCG GGA TCC TCG AGA GTT CAC TCT TGT TGA AG)) and U112-2 (CGC CTC GAG AGA TCT GAT CGT CTG TTA AGA)) and a UNC-112 cDNA plasmid (P13-5) (Qadota et al. 2012) as a template. An amplified cDNA and digested vector (pGAD-C3 with BamHI and BglII) were co-transformed into PJ69-4A yeast strain with pGBDU-PAT-4 with Q308H or I432F (UNC-112 nonbinding mutations) and then transformants were spread onto -His plates. To construct pGBDU-PAT-4 with Q308H or I432F, XhoI fragments of m22 (for Q308H) and m43 (for I432F) were cloned into the pGBDU-C2 XhoI site. A total of 30,000 colonies were screened for suppressors of PAT-4 Q308H. After 3 days of incubation at 30 °C, His+ colonies were 151. Among these 151, 54 clones were isolated and tested for binding to PAT-4 (Q308H). Among 54 isolated clones, 8 clones retained their ability to bind to PAT-4 (Q308H), and the DNA sequences of these clones were determined (Table 2). A total of 846,000 colonies were screened for suppressors of PAT-4 I432F. After 3 days of incubation at 30 °C, His+ colonies were 107. Among 107, 54 clones were isolated and tested for binding to PAT-4 (I432F). Among 54 isolated clones, 14 clones retained their ability to bind to PAT-4 (I432F), and the DNA sequences of these clones were determined (Table 2). To create site-directed mutations of candidate suppressors with single amino acid changes, the following primers were used: for PAT-4 (N37S), TGT CCT TGG AAA TCT TA<u>G</u> TGT GGG AGG ACT CAT GC and GCA TGA GTC CTC CCA CA<u>C</u> TAA GAT TTC CAA GGA CA, for PAT-4 (S185T), CAA TCT TTG CTT CTC AA<u>A</u> CGA ATC TTG ATA TGC GC and GCG CAT ATC AAG ATT CG<u>T</u> TTG AGA AGC AAA GAT TG, for PAT-4 (Q27R), CAC CGA TTT GAA TAT TC<u>G</u> AAG AAG CAT CTC TGT CC and GGA CAG AGA TGC TTC TT<u>C</u> GAA TAT TCA AAT CGG TG, and for PAT-4 (F117S), TTT CTC GGT AAA TTC AT<u>C</u> CAA AGC AAC AAA GAA AT and ATT TCT TTG TTG CTT TG<u>G</u> ATG AAT TTA CCG AGA AA. UNC-112 N-terminal fragments with site-directed mutations were confirmed by DNA sequencing, then cloned into pGAD-C3 and tested for binding to PAT-4 (Q308H) and PAT-4(I432F) by the yeast two hybrid system.

**Table 2.** UNC-112N mutants that restore binding to PAT-4 mutants. Yellow represents single mutations. Blue represents amino acid changes found in two independent clones. Pink represents residues mutated in two independent clones, but those changes result in different amino acid substitutions.

PAT-4 Q308H suppressors
| Mutant | Amino Acid Changes |  |  |  |  |  |  |  |
| --- | --- | --- | --- | --- | --- | --- | --- | --- |
| m22-1-1 | A96V |  |  |  |  |  |  |  |
| m22-4-9 | Q27R | F117S |  |  |  |  |  |  |
| m22-2-33 | V21I | E134D | L362P |  |  |  |  |  |
| m22-4-10 | P214A | Q333E | I338T |  |  |  |  |  |
| m22-3-18 | K118E | S261T | V279E | V325A |  |  |  |  |
| m22-4-17 | K118E | L219S | T233R | V272A |  |  |  |  |
| m22-4-18 | R73Q | N115T | A178T | L330P |  |  |  |  |
| m22-4-4 | S9P | L33F | E44Q | E84D |  |  |  |  |
| m22-1-13 | F117L | E250V | F301L | F315I | Q357H |  |  |  |
| m22-2-34 | L33F | E44Q | G165R | F310S | Y331H |  |  |  |
| m22-6-5 | L33S | V51D | E52D | Q86R | S291P | E371G |  |  |
| m22-4-11 | S29C | Q78E | A96V | S267R | N319S | L393H |  |  |
| m22-3-21 | Q104R | M156L | D232G | H216N | S291P | L330P | L340P | E370D |
| m22-5-34 | contains 1029-1196 only |  |  |  |  |  |  |  |

PAT-4 I432F suppressors
| Mutant | Amino Acid Changes |  |  |  |  |  |  |
| --- | --- | --- | --- | --- | --- | --- | --- |
| m43-4-4 | F301S |  |  |  |  |  |  |
| m43-5-33 | N37S | S185T |  |  |  |  |  |
| m43-6-4 | S116L | Y131C | R293K |  |  |  |  |
| m43-5-19 | V38I | M42L | I284V | F316L |  |  |  |
| m43-3-40 | K118E | S136A | T351P | R395S | S397L |  |  |
| m43-5-35 | S56T | R73Q | Q86L | L187F | S291P | N314D | R395G |

### Co-expression of HA-tagged PAT-4 and myc-tagged UNC-112 in *C. elegans* using heat shock promoters

To create full length UNC-112 with the F301S mutation, a BamHI-BglII fragment of the m43-4-4 clone was inserted into pACT-Q-UNC-112-34 (acceptor plasmid for UNC-112 mutagenesis) (Qadota et al. 2012), resulting in pACT-UNC-112 (F301S). XhoI fragments of UNC-112 cDNA of either wild type (P13-5) or from pACT-UNC-112 (F301S) were cloned into the SalI site of pGBDU-C2, resulting in pGBDU-UNC-112 (wild type or F301S). The SmaI-BglII fragment of pGBDU-UNC-112 (wild type or F301S) was ligated into pBS-myc (kindly provided from Dr. Kozo Kaibuchi, Nagoya University) via the SmaI and BamHI sites, resulting in pBS-myc-UNC-112 (wild type or F301S). To construct pPD49.78- or pPD49.83-myc-UNC-112 plasmids, the SpeI fragment of pBS-myc-UNC-112 (wild type or F301S) was cloned into pPD49.78 or pPD49.83 (gifts from Dr. Andrew Fire, Stanford University) via a NheI site. These vectors contain two different heat shock promoters. pPD49.78/83-HA-PAT-4 (I432F) and pPD49.78/83-myc-UNC-112 (WT or F301S) were mixed with pTG96 (SUR-5::NLS::GFP) as a transformation marker and injected into wild-type N2 worms. Transgenic lines with extrachromosomal arrays containing pPD49.78/83-HA-PAT-4 (I432F)/pPD49.78/83-myc-UNC-112 (WT) (called *sfEx60*) or pPD49.78/83-HA-UNC-PAT-4 (I432F)/pPD49.78/83-myc-UNC-112(F301S) (called *sfEx61*) and pTG96 were established by picking up GFP-positive worms using a GFP dissection microscope. Expression of HA-tagged PAT-4 (I432F) and myc-tagged UNC-112(WT or F301S) proteins was induced by incubation of the transgenic worms at 30 °C for 2 h (heat shock). We prepared Laemmli-soluble protein extracts (Hannak et al. 2002) from these transgenic nematodes with or without heat shock and verified the expression of HA-tagged PAT-4 and myc-tagged UNC-112 proteins by Western blotting, reacting with anti-HA (Sigma-Aldrich H3663, 1/200 dilution) and anti-myc (Sigma-Aldrich H5546, 1/200 dilution).

### Integration of transgenic arrays

The extrachromosomal arrays containing pPD49.78/83-HA-PAT-4 (WT) with pTG96, pPD49.78/83-HA-PAT-4 (I432F) with pTG96, and pPD49.78/83-HA-UNC-PAT-4 (I432F)/pPD49.78/83-myc-UNC-112(F301S) with pTG96 were integrated into the genome by ultraviolet irradiation (Mitani, 1995) with some modifications (P. Barrett, personal communication). The resulting integrated nematode lines are called *sfIs14*, for pPD49.78/83-HA-PAT-4 (WT), *sfIs15*, for pPD49.78/83-HA-PAT-4 (I432F), and *sfIs17*, for pPD49.78/83-HA-UNC-PAT-4 (I432F)/pPD49.78/83-myc-UNC-112(F301S).

### Sustained heat shock overexpression

To investigate the effect of expression of PAT-4 WT, PAT-4 I432F, and PAT-4 I432F/UNC-112 F301S on muscle organization, wild-type worms and worms containing integrated arrasy (sfIs14, sfIs15, and sfIs17) were exposed to 30°C for 24 h. Worms were fixed using the method described below, and immunotained with anti-PAT-6 (to examine M-lines, dense bodies, and cell boundary).

### Immunostaining

Worms were fixed using the method described previously (Nonet et al. 1993; Wilson et al. 2012). Antibody staining with anti-HA (Sigma Aldrich H3663; 1:200 dilution), anti-GFP (Invitrogen A11122, 1:200 dilution), anti-UNC-95 (1:100 dilution; Qadota et al. 2007), and anti-PAT-6 (1:100 dilution; Warner et al. 2013) was performed as described previously (Qadota et al. 2007). Secondary antibodies were anti-rabbit Alexa 488 (Invitrogen) for anti-UNC-95 and anti-GFP, anti-mouse Alexa 594 (Invitrogen) for anti-HA, and anti-rat Alexa 594 (Invitrogen) for anti-PAT-6, each used at 1:200 dilution. Samples were placed on a glass slide with a coverslip containing mounting solution (20 mM Tris (pH 8.0), 0.2 mM DABCO, and 90% glycerol). Images were captured at room temperature with a Zeiss confocal system (LSM510) equipped with an Axiovert 100M microscope and an Apochromat ×63/1.4 numerical aperture oil objective, in ×2.5 zoom mode for Figure 2 and 4, and no zoom mode for Figure 5. The color balances of the images were adjusted by using Adobe Photoshop.

### 3D modeling

For UNC-112 and PAT-4 structure modeling, SWISS-MODEL (Waterhouse et al. 2018) and Phyre2 (Kelley et al. 2015) online tools were used. Human kindlin-3 (7C3M.pdb; Bu et al. 2020), human ILK (3KMW.pdb; Fukuda et al. 2009) and human alpha-parvin (3KMW.pdb; Fukuda et al. 2009) were used as reference crystal structures. The homology model of PAT-4 bound to the CH domain of PAT-6 was based on the human ILK/alpha-parvin complex structure (3KMW.pdb; Fukuda et al. 2009). The Molecular graphics were generated by using Chimera (Pettersen et al. 2004). Single amino acid mutations were inserted using the rotamer tool and then energy minimized in Chimera to minimize interatomic clashes and contacts based on van der Waals radii.

## Data Availability Statement

Worm and yeast strains, and plasmids are available upon request. The authors affirm that all data necessary for confirming the conclusions of the article are present within the article, figures, and tables.

## Results

### Identification of PAT-4 mutants that cannot bind to UNC-112

To gain further insight into the in vivo significance of interaction between PAT-4 (ILK) and UNC-112 (Kindlin), we isolated missense mutations in PAT-4 that result in lack of binding to UNC-112 using the yeast two hybrid system. Using an error-prone PCR approach for generating random mutations, we isolated 16 mutant clones (Table 1). We found that among the 16 clones, 4 clones contain single amino acid changes (N29S, Q308H, I432F, and M464V) (Table 1; Figure 1A). Each mutant PAT-4 protein fails to bind to UNC-112, but can still bind to PAT-6 (α-parvin). Three of these four mutations reside in residues located in the “pseudo kinase” domain, as shown (Q308H, I432F, and M464V). (Figure 1A, Figure 1B). As reported previously, the PAT-4 pseudo kinase domain is essential for binding to UNC-112 (Mackinnon et al. 2002). Since N29S is located outside of the pseudo kinase domain, and there is no crystal structure of ILK or PAT-4 that includes this region, we did not characterize this mutant further. Q308, I432, and M464 are conserved in the human ILK sequence (Supplemental Figure 1). We generated a homology model of the PAT-4 pseudo kinase domain structure based on the crystal structure of the ILK/Parvin complex (Fukuda et al. 2009). According to this homology model, Q308, I432, and M464 are located along the same protein surface (Figure 1C; shown in green). In fact, this surface also contains several residues (I261, F262 and A433) that when mutated permit binding of PAT-4 to UNC-112 D382V (Qadota et al. 2014; shown in red on Figure 1C), and do not overlap with the binding surface of ILK for α-parvin (Fukuda et al. 2009). Q308 is located on an α helix (Figure 1B), and using the program Chimera, upon mutation to Histidine, we found no clashes with neighboring side chains, and this change from polar to a larger polar side chain is predicted to change the local but not the global structure of PAT-4. I432 is located on a loop, and the conservative substitution I432F is predicted to have a minimal effect on either local or global structure using Chimera. M464 is located on the terminal loop, and replacement with valine is also predicted not to affect structure. Since M464 is located C-terminal of the pseudokinase domain (ending with residue 461 as predicted by PFAM), we decided not to characterize it further. Note that position of one non-binding mutation, I432, is next to the amino acid residue A433 that when mutated to serine can restore the binding of PAT-4 to UNC-112 D382V (Qadota et al. 2014; Figure 1C and Figure 6 (with circle)).

**Figure 1.**
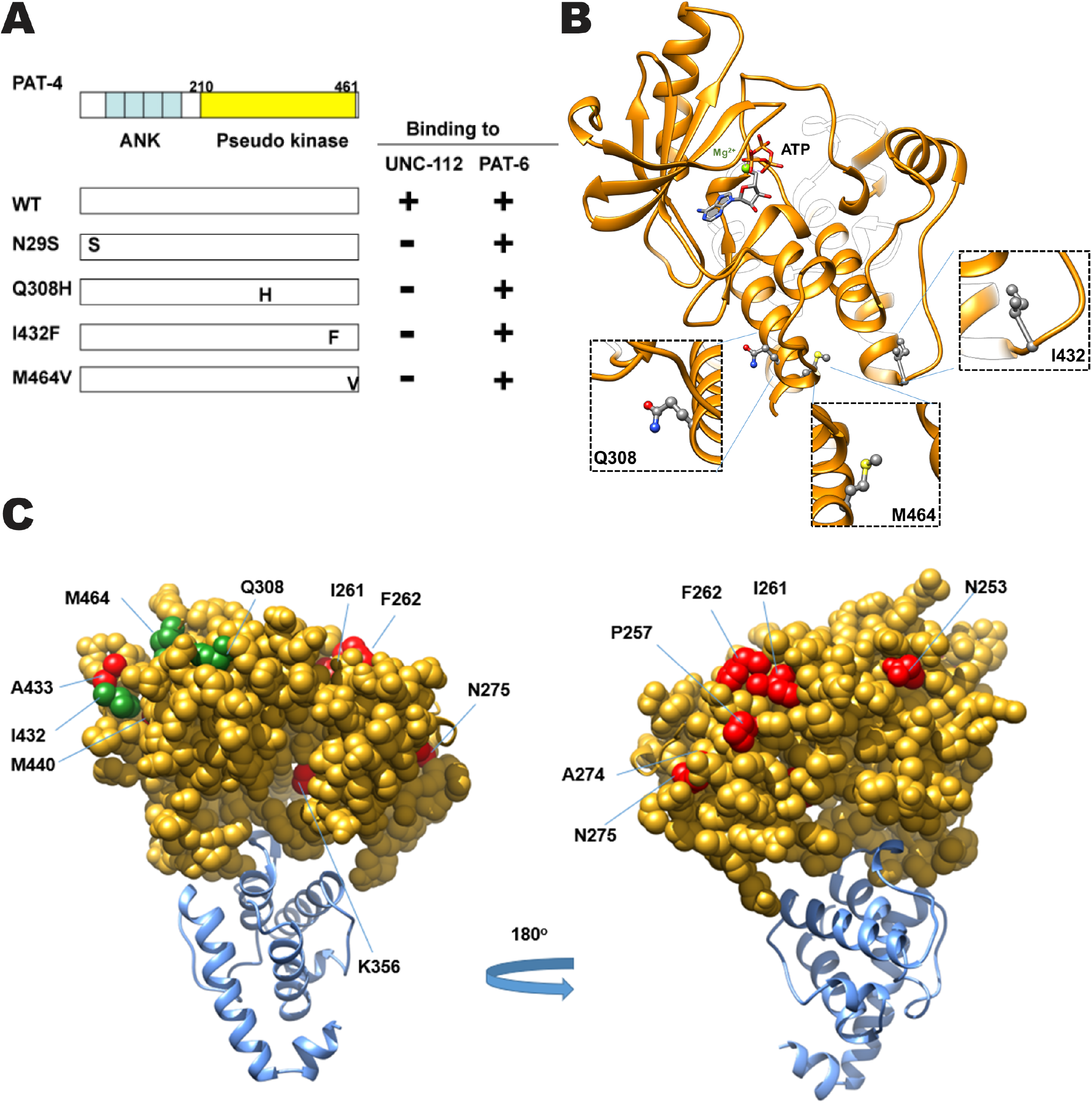
Isolation of PAT-4 missense mutants that fail to interact with UNC-112. A, Schematic representation of domains in PAT-4 (ILK), location of mutations, and results of yeast two hybrid assays. “ANK” (ankyrin) and “Pseudo kinase” are domains predicted by PFAM. Numbers indicate amino acid residue numbers in PAT-4. + represents growth on His- plate and Ade- plate. – represents no growth on either His- plate or Ade- plate. Wild type PAT-4 can bind to UNC- 112 and PAT-6. PAT-4 with N29S, Q308H, I432F, or M464V cannot bind to UNC-112, but still can bind to PAT-6. B, Homology model of PAT-4 structure based on the crystal structure of human ILK (PDB: 3KMW) (Fukuda et al. 2009) using Swiss-Model (Waterhouse et al. 2018), and showing the residues in the pseudo kinase domain that are mutated. The color scheme matches the colors of the domains in the schematic of part A. Three amino acid changes appearing in part A (Q308H, I432F, M464V) are shown in enlarged windows. C, Homology model of PAT-4 bound to the CH domain of PAT-6 based on the human ILK/α-parvin complex structure (3KMW.pdb; Fukuda et al. 2009) modelled with SWISS-MODEL (Waterhouse et al. 2018). PAT-4 kinase domain is shown in yellow in the space filling mode, the CH domain of PAT-6 is shown in dark cyan. The residues mutated in the 3 PAT-4 mutants identified here that fail to bind to UNC-112 (Q308H, I432F, M464V) are indicated in green. Residues that when mutated permit binding of PAT-4 to UNC-112 D382V (Qadota et al. 2014) are indicated in red. All of these residues mutated in PAT-4 are on a surface that is not covered by or does not overlap with the binding site for PAT-6 (α-parvin). This surface appears to have two clusters: one side with residues M440, I432, A433, M464 and Q308, and the other side with residues N275, A274, P257, F262 and I261.

### PAT-4 mutant proteins that cannot bind to UNC-112 do not localize to muscle IACs

To examine whether the Q308H or I432F mutations affect the localization of PAT-4 in *C. elegans* muscle, we created transgenic animals expressing HA tagged wild type or mutant PAT-4 proteins. Expression of HA tagged PAT-4 (WT, Q308H, and I432F) was confirmed by western blot, and was strictly controlled by the heat shock promoters of our constructs (Figure 2A). After heat shock, transgenic animals were immunostained with anti-HA (to localize HA tagged PAT-4) and anti-UNC-95 (to identify muscle integrin adhesion complex structures). HA-PAT-4 WT localized to dense bodies and M-lines similar to UNC-95, but HA-PAT-4 Q308H and HA-PAT-4 I432F did not localize to dense bodies and M-lines (Figure 2B), suggesting that PAT-4 localization at muscle attachment structures requires binding to UNC-112 and since these animals express endogenous levels of PAT-6 (α-parvin), the interaction of PAT-4(ILK) to PAT-6 (α-parvin) is not sufficient for proper localization of PAT-4 (ILK).

**Figure 2.**
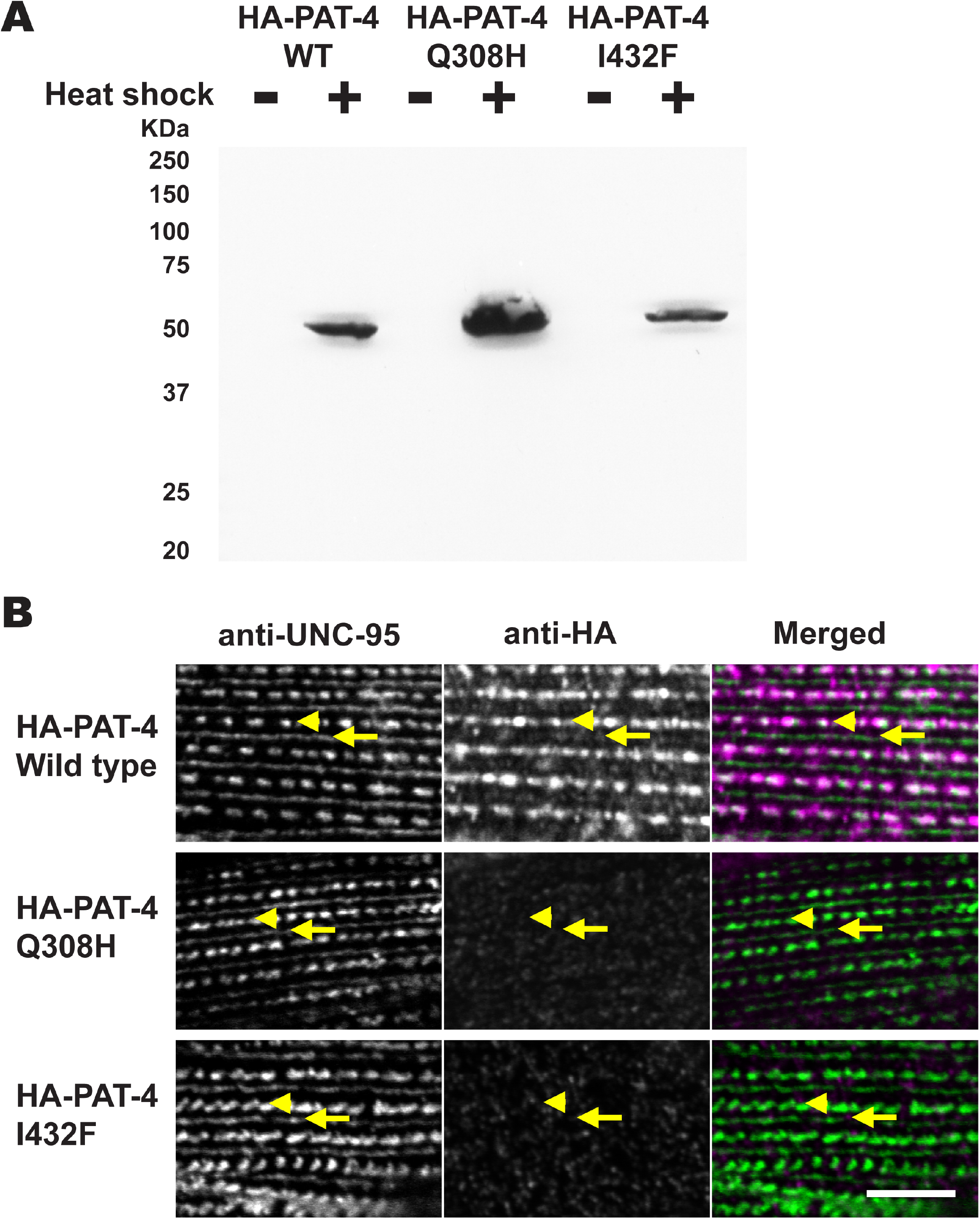
PAT-4 Q308H and PAT-4 I432F mutant proteins that cannot bind to UNC-112 do not localize to integrin adhesion complexes. A, Western blot of lysates from transgenic nematodes carrying HA-tagged PAT-4 wild type, Q308H or I432F, expressed from a heat shock promoter, with heat shock (+) or without heat shock (-), reacted with anti-HA. Note that the apparently different level of expression of HA-PAT-4 is due to the variable rate of transmission of these extrachromosomal arrays in the different transgenic lines. B, localization of heat shock-expressed HA-tagged PAT-4 wild type, Q308H, and I432F in transgenic animals. Worms were immunostained with anti-HA to detect the transgenic PAT-4 and with anti-UNC-95 to visualize the optical plane in body wall muscle cells that contain the integrin adhesion complexes (dense bodies and M-lines). Wild type HA-PAT-4 localizes normally to dense bodies and M-lines. However, Q308H and I432F HA-PAT-4 fail to localize to these structures. White bar, 10 μm.

### Identification of UNC-112 mutants that can restore binding to PAT-4 (Q308H) or PAT-4 (I432F)

To confirm the possibility that UNC-112 binding is sufficient for the proper localization of PAT-4 to muscle IACs, we isolated UNC-112 suppressor mutations that restore the ability of PAT-4 Q308H or I432F mutant proteins to interact with UNC-112. From suppressor screening of PAT-4 Q308H, we isolated 6 suppressor mutant UNC-112 N-terminal clones. From suppressor screening of PAT-4 I432F, we isolated 14 suppressor mutant UNC-112 N-terminal clones (Table 2). Among the total of 20 suppressor mutant clones, there were two single amino acid changes, F301S and A96V, (Table 2; Figure 3A). Among remaining 18 mutant clones, there were two clones containing two mutations, m43-5-33 (N37S and S185T) and m22-4-9 (Q27R and F117S). To identify which single amino acid changes are critical for suppression, we created each single mutation by site-directed mutagenesis, then tested for PAT-4 Q308H or I432F binding using the yeast two hybrid assay. As a result, an UNC-112 N-terminal half with N37S showed binding to PAT-4 I432F, but UNC-112 N-terminal halves with the three other single mutations did not show binding to PAT-4 Q308H or I432F. In the case of m43-5-33, since UNC-112N with N37S showed binding to PAT-4 I432F, S185T is not essential for suppression. In the case of m22-4-9, since UNC-112N with either Q27R or F117S did not show binding to PAT-4 Q308H, we concluded that two mutations (Q27R and F117S) are required for binding to PAT-4 Q308H. Taken together, we identified four sets of suppressor mutations; F301S, N37S, A96V, and Q27R plus F117S (Figure 3A). Furthermore, all four mutation sets (F301S, N37S, A96V, and Q27R plus F117S) can bind to both PAT-4 Q308H and PAT-4 I432F. Previously (Qadota et al. 2020), we reported generating a homology model of UNC-112 based on the crystal structure of human kindlin-3 (Bu et al. 2020). We placed our suppressor mutations on this model. In this model (Figure 3B), N37, A96, and F301 are located along one side of UNC-112, fairly close together. N37 is located on a loop and the conservative substitution to serine, is predicted by Chimera to have minimal effect on local or global structure. Similarly, A96 is also located on a loop and the conservative substitution to valine is predicted to have minimal effect on the local or global structure of UNC-112. F301 is also located on a loop, and changing the large hydrophobic residue phenylalanine to the small polar residue serine is predicted to to change the local but not global structure of UNC-112. Taken together, all three single amino acid changes seem to have no or minimal effect on the global structure of UNC-112, and are suggested to be involved in PAT-4 binding. Interestingly, the position of one suppressor mutation, F301, is next to the amino acid residue found in the PAT-4 non-binding mutation, E302G (Qadota et al. 2021; Figure 3C; Figure 6 (with rectangle)), further highlighting the importance of the region F301-E302 in UNC-112 for PAT-4 binding.

**Figure 3.**
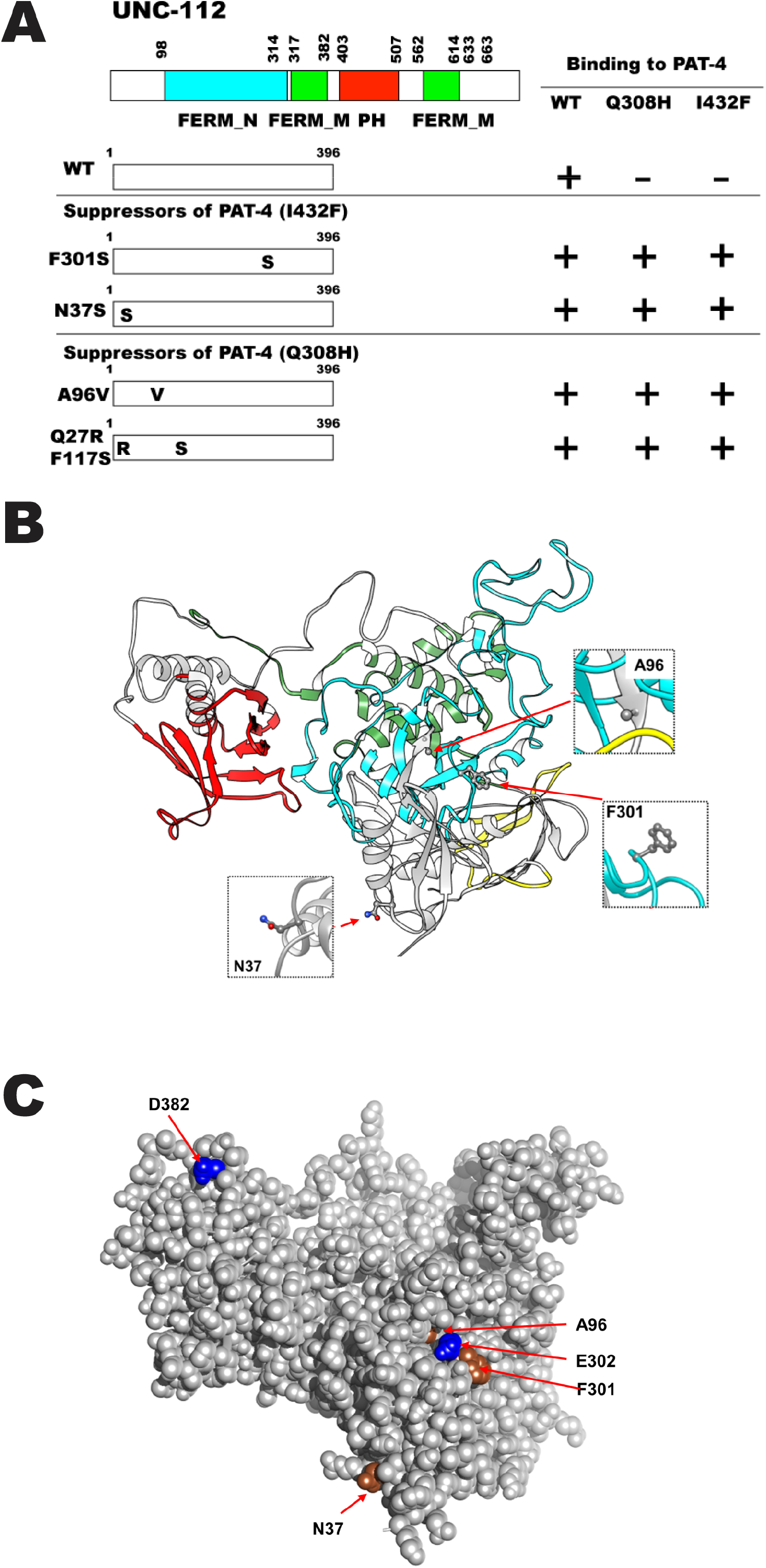
Location of amino acid changes in UNC-112 suppressor mutant proteins that restore binding to PAT-4 (Q308H) and PAT-4 (I432F) A, Schematic representation of domains in UNC-112 (kindlin), location of mutations, and results of yeast two hybrid assays. FERM_N, FERM_M, and PH are domains predicted by PFAM. Numbers indicate amino acid residue numbers in UNC-112. + represents growth on His- plate and Ade- plate. – represents no growth on either His- plate or Ade- plate. Wild type UNC-112 N-terminal half cannot bind to PAT-4 containing either Q308H or I432F mutations. UNC-112 N-terminal half with F301S, N37S, A96V, or Q27Q/F117S can bind to PAT-4 with Q308H and I432F. B, Structure of UNC-112 (Qadota et al. 2020) based on the human kindlin-3 3D structure (PDB: 7C3M) (Bu et al. 2020) modelled with Swiss-Model (Waterhouse et al. 2018), and showing corresponding mutated residues. The color scheme matches the colors of the domains in the schematic of part A. Residues that were affected by single amino acid changes appearing in part A (N37, A96, and F301) are shown in enlarged windows. C, Spacefilling model of UNC-112 highlighting the location of the N37, A96, and F301 (in brown) identified in this study that when mutated restore the ability of PAT-4 Q308H and I432F mutants to bind to UNC-112, and D382 and E302 residues (in blue) that when mutated fail to bind to PAT-4 (Qadota et al. 2012; Qdota et al. 2021).

### UNC-112 F301S can restore the ability of PAT-4 I432F to localize to M-lines and dense bodies in muscle

To examine whether UNC-112 F301S can restore the localization of PAT-4 I432F to muscle IACs, we created two transgenic lines: (i) co-expression in body wall muscle of HA-PAT-4 I432F and myc-UNC-112 wild type, and (ii) co-expression in body wall muscle of HA-PAT-4 I432F and myc-UNC-112 F301S. Heat shock induced expression of each protein in the two transgenic lines was demonstrated by western blot using antibodies to HA and myc (Figure 4A). We co-immunostained with anti-HA (to localize HA-PAT-4) and with anti-UNC-95 (to mark the IACs at M-lines and dense bodies). As indicated in Figure 4B, it is clear that although myc-UNC-112 wild type fails to localize HA-PAT-4 I432F to IACs (similar to results in Figure 2B), myc-UNC112 F301S can permit at least partial localization of HA-PAT-4 I432F to IACs in muscle.

**Figure 4.**
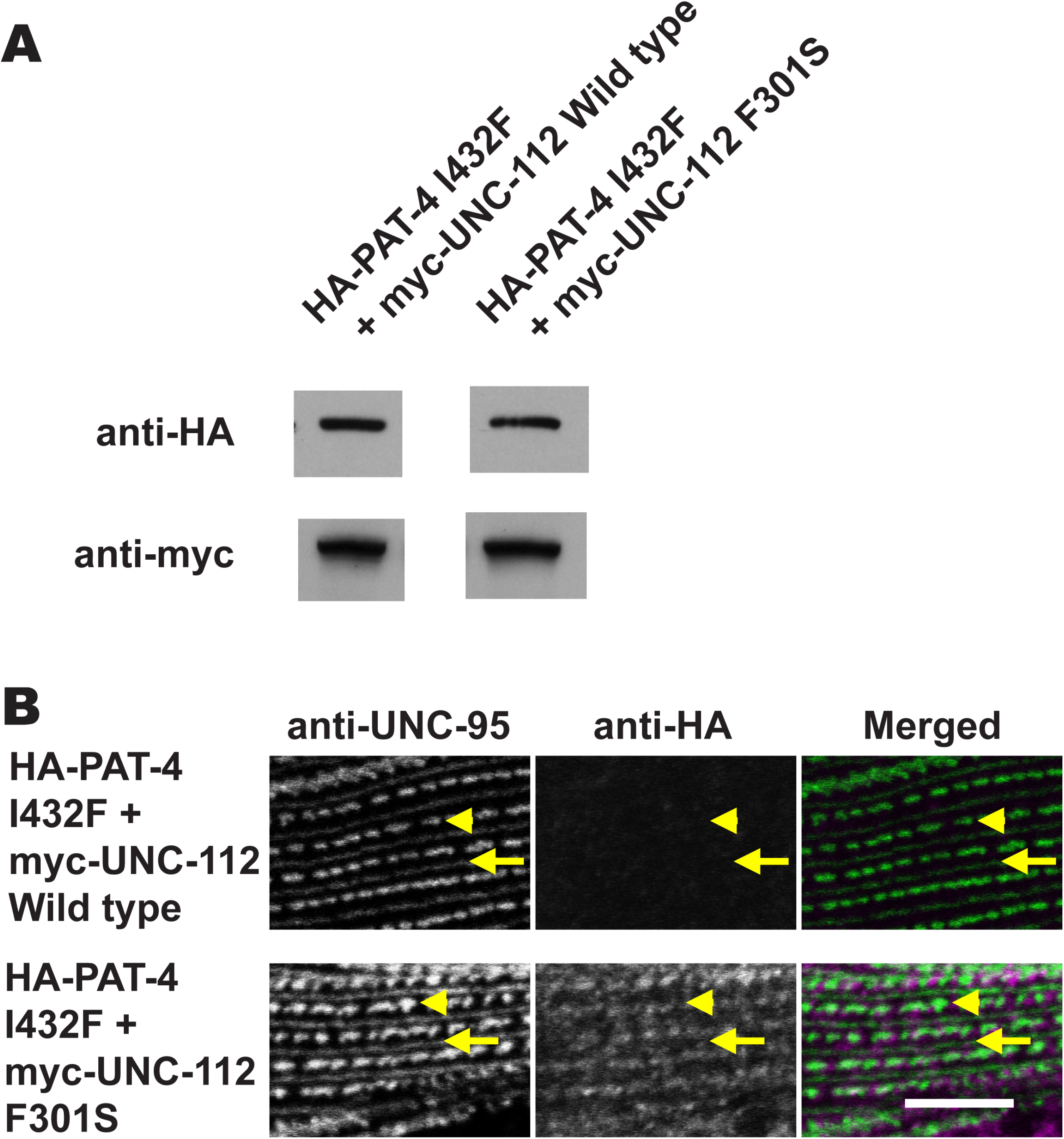
Co-expression of HA-PAT-4 I432F and myc-UNC-112 F301S restores ability of PAT-4 I432F to localize to integrin adhesion sites. A, Western blot of lysates from transgenic worms expressing from a heat shock promoter HA-tagged PAT-4 I432F and either myc-tagged UNC-112 wild-type (WT) or UNC-112 F301S. Lysates were prepared after heat shock (+) and reacted with anti-HA or anti-myc antibodies. Expression of the HA- and myc-tagged proteins depended on heat shock. B, Localization of heat shock-expressed HA-tagged PAT-4 I432F in the presence of co-expressed myc-tagged UNC-112 wild type, or myc-tagged UNC-112 F301S. Adult worms were immunostained with anti-HA to detect transgenic PAT-4 and with anti-UNC-95 to visualize the optical plane of body wall muscle cells that contain dense bodies and M-lines (integrin adhesion sites). HA-PAT-4 I432F fails to localize to these structures in the presence of myc-UNC-112 WT but does localize to these structures in the presence of myc-UNC-112 F301S. Yellow arrowheads mark dense bodies, whereas yellow arrows mark M-lines. White bar, 10 μm.

### Overexpression of PAT-4 I432F results in disorganization of adhesion complexes at muscle cell boundaries and this disorganization is suppressed by co-expression of UNC-112 F301S

The PAT-4 I432F mutant protein cannot bind to UNC-112, but can bind to PAT-6 (Figure 1). In addition, the PAT-4 I432F mutant protein cannot localize to any of the integrin adhesion complexes in nematode muscle, including M-lines and dense bodies (Figure 2), or adhesion plaques at muscle cell boundaries (data not shown). We reasoned that if an excessive amount of PAT-4 I432F was expressed, PAT-4 I432F could bind to PAT-6, but because this PAT-4 I432F cannot bind to UNC-112, it would not localize to integrin adhesion complexes, and thus the amount of PAT-6 available to localize to integrin adhesion complexes would be reduced. We tested this idea by heat shock overexpression of PAT-4 WT, PAT-4 I432F, and PAT-4 I432F/UNC-112 F301S. Young adult worms were exposed to 30°C for 24 hours. In wild type worms, this heat shock treatment had no effect on the localization of PAT-6 to any of the integrin adhesion complexes (M-lines, dense bodies or adhesion plaques at muscle cell boundaries; labeled wild type in Figure 5). Overexpression of PAT-4 WT also had no effect on muscle attachment structures (labeled OE: PAT-4 WT in Figure 5). However, overexpression of PAT-4 I432F resulted in mild to moderate disorganization of dense bodies, and severe disorganization of the muscle cell boundaries (OE: PAT-4 I432F in Figure 5). In particular, the adhesion plaques at the cell boundaires are more loosely organized than in wild type, and there appear to larger gaps between adjacent muscle cells. The defect in cell boundaries caused by overexpression of PAT-4 I432F could be largely suppressed by co-expression of UNC-112 F301S (OE: PAT-4 I432F + UNC-112 F301S in Figure 5).

**Figure 5.**
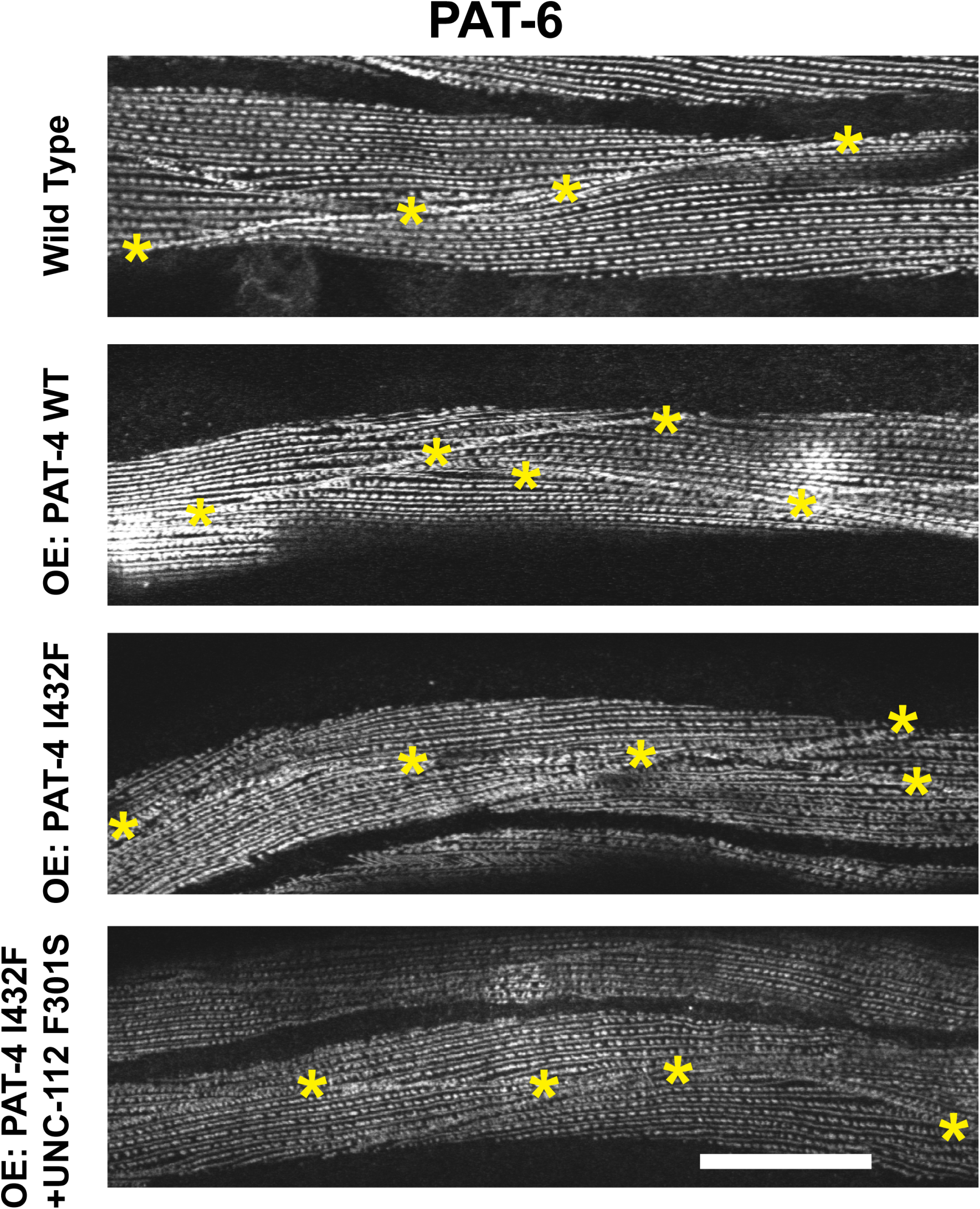
Sustained overexpression of PAT-4 I432F results in disorganization of muscle cell boundary integrin adhesion complexes and this disorganization is suppressed by co-expression of UNC-112 F301S. Localization of PAT-6 (α-parvin) at M-lines, dense bodies, and muscle cell boundaries in wild type worms, and worms carrying integrated arrays of PAT-4 WT, PAT-4 I432F, and PAT-4 I432F + UNC-112 F301S. Young adult worms were exposed to 30°C 24 hours, then fixed and immunostained with anti-PAT-6. Yellow asterisks mark muscle cell boundaries. Overexpression of PAT-4 I432F results in reduced accumulation of PAT-6 at the boundaries, abnormal space between adjacent muscle cells, and a mild disruption of the structure of dense bodies (appearing as a series of “Us” rather than dashes). White bar, 10 μm.

## Discussion

### PAT-4 (ILK) protein surface that interacts with UNC-112(Kindlin)

The goal of this study was to obtain more information about an interacting surface between PAT-4 and UNC-112. Previously, we identified extragenic suppressor mutations in PAT-4 that restore binding to UNC-112 D382V (see Figure 6 in which D382V (Qadota et al. 2012) is in blue font for UNC-112, and PAT-4 suppressors (Qadota et al. 2014) are in red font for PAT-4). In a homology model of PAT-4, based on the ILK-parvin co-crystal structure (Fukuda et al. 2009), we had reported that residues mutated in the suppressors are located on one side of the PAT-4 protein, different from the surface of ILK (PAT-4) that binds to α-parvin (PAT-6) (Qadota et al. 2014). The three PAT-4 mutations that we have described here (see Figure 6 in which Q308H, I432F, and M464V are in green font) that cannot bind to UNC-112, are also located on the side of PAT-4 as are the nine extragenic suppressor mutations (Figure 1C). Thus, all of the residues mutated in PAT-4 that fail to bind to UNC-112 or when mutated can restore binding to UNC-112 D382V are on a surface that is not covered by or does not overlap with the binding site for PAT-6 (α-parvin). Furthermore, this surface appears to have two clusters: one side with residues M440, I432, A433, M464 and Q308, and the other side with residues N275, A274, P257, F262 and I261. We hypothesize that these two clusters are crucial for interaction with UNC-112 (kindlin).

As we noted in Results, the I432F mutation results in non-binding to UNC-112, whereas the next amino acid residue A433 when mutated to serine, restores binding to UNC-112 D382V (black rectangles for PAT-4 in Figure 6), further suggesting that the surface of PAT-4 containing non-binding and suppressor mutations is likely to be involved in binding to UNC-112. In the case of human ILK, since kindlin-2 L357 lies along the non-polar face of an amphipathic helix important for ILK binding (Fukuda et al. 2014; Huet-Calderwood et al. 2014), it was hypothesized that a hydrophobic patch on ILK might be important for kindlin binding. Guided by this hypothesisi, I427 of ILK was found to be essential for kindlin binding. Additional surface-exposed residues near I427 were tested for kindlin binding, and one of them, K423, was also found to be important for kindlin binding (Kadry et al. 2018). Corresponding to amino acid residues K423 and I427 in human ILK, are R438 and I442 in worm PAT-4. We should note that R438 and I442 are nearby one of the PAT-4 mutations described here that fail to bind to UNC-112, namely I432. For human ILK, the region corresponding to Q308 in PAT-4, has not yet been reported to be characterized.

**Figure 6.**
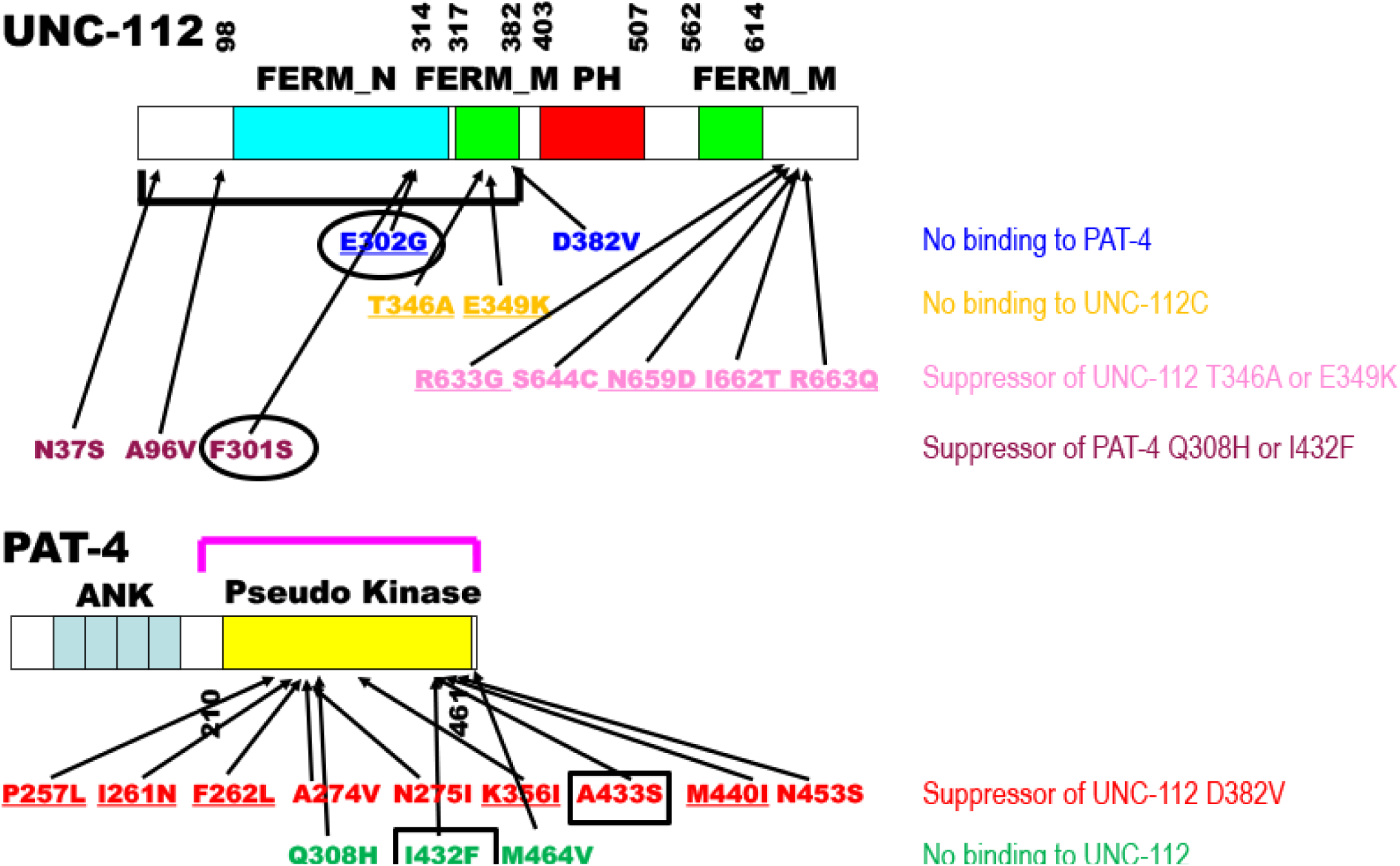
Summary of single amino acid substitutions in UNC-112 and PAT-4 that affect intermolecular interaction between UNC-112 and PAT-4, and intramolecular interaction between the N- and C-terminal halves of UNC-112. Schematic representation of domains in UNC-112 (kindlin) and PAT-4 (ILK), location of mutations, and results of yeast two hybrid assays. “FERM_N”, “FERM_M”, “PH”, “ANK”, and “Pseudo kinase” are domains predicted by PFAM. Numbers indicate amino acid residue numbers in UNC-112 and PAT-4. The black bracket represents the region of UNC-112 that binds to PAT-4 (Mackinnon et al. 2002). The purple bracket represents the region of PAT-4 that binds to UNC-112 (Mackinnon et al. 2002). Blue amino acid changes in UNC-112 (D382V and E302G) are UNC-112 mutants that cannot bind to PAT-4 (Qadota et al. 2012; Qdota et al. 2021). Orange amino acid changes are UNC-112 N-terminal mutants (T346A and E349K) that cannot bind to the UNC-112 C-terminal half (Qadota et al. 2012). Pink amino acid changes (R633G, S644C, N659D, I662T, and R663Q) restore the ability of UNC-112 N-terminal mutant proteins (T346A and E349K) to bind to the UNC-112 C-terminal half (Qadota et al. 2020). Purple amino acid changes in UNC-112 (N37S, A96V, and F301S) restore the ability of UNC-112 to bind to PAT-4 Q308H or I432F (this study). Red amino acid changes in PAT-4 (P257L, I261N, F262L, A274V, N275I, K366I, A433S, M440I, N453S) restore the ability of PAT-4 to bind to UNC-112 D382V (Qadota et al. 2014). Green amino acid changes in PAT-4 (Q308H, I432F, and M464V) result in PAT-4 that cannot bind to UNC-112 (this study). Underlining represents conserved amino acid residues (UNC-112 for kindlin, and PAT-4 for ILK). Black circles in UNC-112 (around E302G and F301S) and black rectangles in PAT-4 (around A433S and I432F) indicate that a non-binding mutation and a suppressor mutation are located in consecutive amino acid residues.

### UNC-112 (Kindlin) protein surface that interacts with PAT-4 (ILK)

We have reported previously that the UNC-112 N-terminal half containing the FERM_N and part of the FERM_M domains binds to PAT-4, and that a missense mutation in the FERM_M domain (D382V) abolishes binding to PAT-4 (Qadota et al. 2012). For mammalian kindlin-2, it has been reported that the N-terminal half of FERM_M domain is responsible for binding to ILK, and that single amino acid changes in this region abolish binding to ILK (Fukuda et al. 2014). Recently, we reported that a missense mutation in the FERM_N domain of UNC-112, E302G, also results in diminished binding to PAT-4(ILK) (Qadota et al. 2021), thus providing additional evidence that the FERM_N domain is critical for binding to PAT-4(ILK). In the study reported here, we found that the mutations in UNC-112 that restore binding to PAT-4 Q308H or I432F, are also located in the FERM-N domain (F301), or N-terminal of the FERM-N domain (N37S and A96V)(in purple font for UNC-112 in Figure 6). In fact, all three residues affected are located along the same side of UNC-112 in our homology model of UNC-112 (Figure 3B). It is interesting to note that one of the suppressor mutations, F301S, resides in an amino acid next to E302, that when mutated to glycine cannot bind to PAT-4 (Qadota et al. 2021). The previous data and data shown here expand the region and the residues of kindlin family proteins that are likely involved in the interaction with ILK.

### PAT-4 binding to UNC-112 is required for the localization of PAT-4 to IACs

Loss of function mutations in *pat-4* and *unc-112* are Pat/embryonic lethal (Mackinnon et al. 2002; Rogalski et al. 2000), because a functioning myofilament lattice is not formed in embryonic muscle. Based on examining the localization of proteins in embryonic muscle of dying *pat-4* and *unc-112* embryos, it has been demonstrated that PAT-4 is required for proper localization of UNC-112, and that UNC-112 is required for proper localization of PAT-4 (Mackinnon et al. 2002). In adult *C. elegans* muscle cells, by expressing an HA-tagged UNC-112 mutant (D382V) protein that cannot bind to PAT-4, it was shown that PAT-4 binding to UNC-112 is required for UNC-112 to localize to integrin adhesion complexes (M-lines and dense bodies) (Qadota et al. 2012). In this study, we isolated PAT-4 mutants that cannot bind to UNC-112, and expressed those two mutant PAT-4 proteins in adult *C. elegans* muscles, and found that UNC-112 binding is required for PAT-4 localization to integrin adhesion complexes (Figure 2). Furthermore, we found that co-expression of a mutant (F301S) UNC-112 protein that restores binding to mutant PAT-4(I432F), also restores the localization of the mutant PAT-4 to integrin adhesion complexes (Figure 4). Similar results have been reported for human ILK. Human ILK with either K423D or I427E in the pseudo kinase domain (Supplemental Figure 1) could not localize to focal adhesions of CHO cells (Kadry et al. 2018). Interestingly, both the PAT-4 mutants and ILK mutants did not show an obvious defect in protein stability (Figure 2; Kadry et al. 2018)). Our PAT-4 mutant proteins cannot bind to UNC-112, but still can bind to PAT-6 (α-parvin) (Figure 1), indicating that the overall structure and stability of the PAT-4 mutant proteins are intact. Moreover, since our four PAT-4 mutant proteins have amino acid changes in the pseudo kinase domain (Q308H, I432F, M464V), or in the N-terminus (N29S), and not in the ankyrin repeat region which is known to bind to UNC-97 (PINCH) (Mackinnon et al. 2002; Norman et al. 2007), suggesting that these PAT-4 mutant proteins, when co-expressed with suppressor mutant UNC-112 (F301S) protein can still form a complete four protein complex (UNC-112/PAT-4/PAT-6/UNC-97) in vivo. Since we previously demonstrated by co-pelleting from worm extracts that UNC-112(Kindlin), PAT-4(ILK), PAT-6(α-parvin) and UNC-97(PINCH) form a tight complex (Qadota et al. 2014), we propose the name, “KIPP complex”, for this four-protein complex.

### Integrin adhesion complexes at muscle cell boundaries are more dynamic than M-lines and dense bodies

*C. elegans* muscle cells attach to extracellular matrix through three structures: M-lines, dense bodies, and adhesion plaques at muscle cell boundaries (Qadota et al., 2017). These three attachment structures contain different sets of proteins, but a number of proteins are in common, including PAT-3 (β-integrin), UNC-112 (Kindlin), PAT-4 (ILK), PAT-6 (α-parvin), UNC-97 (PINCH), and UNC-95, that are located near the muscle cell membrane. In this study, we reported that overexpression of PAT-4 I432F, that cannot bind to UNC-112, results in disorganization of muscle cell boundaries and mild disorganization of dense bodies (Figure 5). We interpret these results in the following way: PAT-4 I432F binds to PAT-6, but this PAT-4/PAT-6 complex cannot localize to attachment structures due to PAT-4 I432F’s inability to bind to UNC-112, and as a result, the amount of PAT-6 available to localize to adhesion complexes is reduced. The reduced level of PAT-6 results in less downstream proteins at these structures. In fact, although the null state for *pat-6* is Pat embryonic lethal (Lin et al., 2003), *pat-6* RNAi beginning from the L1 larval stage results in disorganization of adult myofibrils (unpublished data). In addition, the effect of PAT-4 I432F overexpression was limited to muscle cell boundaries and dense bodies (somewhat), but not to M-lines. Our previous study suggests that the adhesion plaques at muscle cell boundaries are more dynamic than dense bodies or M-lines (Moody et al. 2020): PIX-1 is a Rac GEF that is localized to all three integrin adhesion structures (M-lines, dense bodies and boundaries), but is only required at muscle cell boundaries. Either loss of function or overexpression of PIX-1 results in the same cell boundary defect (Moody et al. 2020), suggesting that cell boundary structures require rapid turn over of component proteins, and is more dynamic than M-lines and dense bodies. In the future, we might be able to test this hypothesis by measuring and comparing protein exchange rates at the cell boundary, dense bodies and M-lines by, for example, FRAP analysis.

### Genetic analysis of the UNC-112 to PAT-4 interaction

Figure 6 summarizes our results to date using a random mutagenesis approach (Qadota and Benian 2014) to isolate non-binding mutants and their suppressor mutants (Qadota et al. 2012; Qadota et al. 2014, Qadota et al. 2020; Qadota et al. 2021; this study). Other than deletion and non-sense mutations, there are reports about two *unc-112* alleles (no reports for *pat-4*) : First, the Unc allele, *unc-112(r367)*, is a missense mutation, T85I (Rogalski et al. 2000). The molecular nature of UNC-112 T85I has not been examined, and thus, we do not know if the phenotype is due to modifed binding to PAT-4 or PAT-3, or whether it results in a unstable protein. Second, there is one unusual allele created by CRISPR/Cas9, *unc-112(kq715)*, L715E, near the end of protein, which shows a defect in the migration of the distal tip cell (Park et al. 2020), but whether there was an effect on the attachment structures of body wall muscle, was not reported. Using this accumulated set of mutations that affect interaction between UNC-112 and PAT-4, we can roughly estimate the binding surface between these two proteins, and key residues of each protein responsible for this interaction. However, to clarify the binding surface definitively, analysis of a co-crystal structure of UNC-112 (Kindlin) and PAT-4 (ILK) will be required, and so far, no such co-crystal structure for any kindlin/ILK pair has been reported. Hopefully, our mutational analysis will be of some use to validate co-crystal structures. Moreover, we have not yet explored the in vivo phenotype of the mutations in *unc-112* and *pat-4* shown in Figure 6, but we showed reduced localization of PAT-4 and UNC-112 proteins and the deliterious effects of mutant PAT-4 and UNC-112 proteins in adult muscle (this study; Qadota, et al. 2012). Since both UNC-112 and PAT-4 require each other (Mackinnon et al. 2002), individual homozygous mutations might result in Pat/embryonic lethality, but in the presence of a suppressor mutation, the worms might bypass the embryonic lethality and either display an adult Unc phenotype or appear like wild type. In the future, using CRISPR/Cas9 technology, we hope to create homozygous non-binding mutants with or without homozygous suppressor mutations, and then examine for embryonic and adult stage muscle defects. Through this type of in vivo genetic analysis, we would be able to reveal the in vivo meaning of the intertaction of UNC-112 and PAT-4 in the excellent genetic model, *C. elegans*.

## Acknowledgments

We thank Dr. Andrew Fire (Stanford University) for *C. elegans* expression vectors (pPD49.78 and pPD49.83) and Dr. Kozo Kaibuchi (Nagoya University) for pKS-HA and pBS-myc. The *C. elegans* wild type strain, N2, was provided by the Caenorhabditis Genetics Center, which is funded by the NIH Office of Research Infrastructure Programs (P40 OD010440).

## Funding and additional information

This study was supported in-part, by a previous grant from the American Heart Association (11GRNT7820000), and a more recent grant from NIH (1R01HL160693).

## Figures and figure legends

**Supplemental Figure 1**. Alignment of human ILK and worm PAT-4. The blue stars indicate residues in the PAT-4 pseudo kinase domain that when mutated fail to bind to UNC-112.

